# Supervised and unsupervised deep learning-based approaches for studying DNA replication spatiotemporal dynamics

**DOI:** 10.1101/2024.05.09.593366

**Authors:** Julian Ng-Kee-Kwong, Ben Philps, Fiona N. C. Smith, Aleksandra Sobieska, Naiming Chen, Constance Alabert, Hakan Bilen, Sara C. B. Buonomo

## Abstract

In eukaryotic cells, DNA replication is organised both spatially and temporally, as evidenced by the stage-specific spatial distribution of replication foci in the nucleus. Despite the genetic association of aberrant DNA replication with numerous human diseases, the labour-intensive methods employed to study DNA replication have hindered large-scale analyses of its roles in pathological processes. In this study, we first demonstrate that a convolutional neural network trained to classify S-phase stages based on DAPI and EdU patterns could identify altered replication dynamics in *Rif1*-deficient mouse embryonic stem cells (mESCs), revealing a skewed distribution across the various S-phase stages. Given the possible practical limitations associated with a supervised framework, we proceed to show that the abnormal replication profile of *Rif1*-deficient mESCs could further be detected by an unsupervised approach (based on self-supervised representation learning), which could additionally reconstruct progression through S-phase. Finally, we extend our approach to a well-characterised cellular model of inducible deregulated origin firing, involving cyclin E overexpression. Through parallel EdU- and PCNA-based analyses, we demonstrate the potential applicability of our method to patient samples, offering a means to identify the contribution of deregulated DNA replication to a plethora of pathogenic processes.

## Introduction

DNA replication in eukaryotes follows a defined chronological order, known as the replication-timing (RT) program^1^. The importance of the RT program is supported by its conservation throughout eukaryotic evolution and by its robust control—perturbing the RT program has indeed proven extremely difficult^2^. This temporal control coincides with the spatial organisation of DNA replication within the nucleus^1,3^. For a given genomic region, the timepoint at which it is replicated in S-phase is generally highly correlated with its localisation in the nucleus. Such an association is also evident globally, where a broad correspondence has been demonstrated between early- and late-replicating chromatin, identified by Repli-seq^4^, and the A and B compartments, identified by Hi-C analyses^5–7^.

As large genome segments are replicated, clusters of synchronously firing replication origins manifest as distinct foci in the nucleus^8,9^. Visualisation of replication foci by fluorescence microscopy is made possible by labelling actively replicating DNA with nucleoside analogues, such as 5-bromo-2’-deoxyuridine (BrdU) or 5-ethynyl-2ʹ-deoxyuridine (EdU), followed, respectively, by immunofluorescence or click chemistry with fluorescent azides^10^. Alternatively, immunostaining for fork-associated proteins such as proliferating cell nuclear antigen (PCNA) gives comparable results^11–13^. These assays have revealed consecutive and distinct spatial patterns of replication fork distribution, representing the temporal progression through early, mid and late S-phase^14^. Subsequent studies extended this classification to five or six observable patterns^8^, consistent across various cell lines^15^. In mouse embryonic stem cells (mESCs)^16^, up to six patterns have been classified: S1 to S6^17^.

Aberrant DNA replication is linked to several diseases^18,19^. For example, the deregulated firing of origins of replication caused by the upregulation of oncogenes is an early step of cell transformation^20^. The genomic methods commonly employed to study deregulated DNA replication, which are based on BrdU or EdU pulses, are not easily applicable to human studies and are also expensive. PCNA immunostaining on tissue cryosections, combined with high-throughput imaging analysis would enable the study of patient samples, addressing the question of how early during pathogenesis deregulation of DNA replication dynamics takes place and also how commonly it occurs. However, the analysis of S-phase progression by manual annotation of DNA replication patterns is subjective, laborious and time-consuming, particularly when performed at scale. A standardised, unbiased and scalable method to study DNA replication progression would therefore be an important advancement. High-throughput image acquisition combined with machine learning provides a promising avenue towards achieving this.

Machine learning-based approaches have previously been employed to classify the broader cell cycle stages (G1, S, G2 and M) and the four phases of mitosis (prophase, metaphase, anaphase and telophase)^21–24^. To our knowledge, however, the application of convolutional neural networks specifically to S-phase classification has not been attempted and could obviate the need to rely on 4ʹ,6-diamidino-2-phenylindole (DAPI) intensity profiles—which can be difficult to standardise across experiments—or tedious manual categorisation of individual nuclei.

Here, we first evaluate, in a supervised setting, the performance of a convolutional neural network in classifying the stages of S-phase based on DAPI and EdU patterns. Through visualisation of the model-generated image embeddings, we demonstrate that the trained model can reconstruct wild-type S-phase progression based on categorical labels, consistent with past findings where this was attempted for cell cycle progression^21^. We then apply the trained model to the analysis of S-phase progression in *Rif1*-deficient cell lines. RIF1 is an adaptor of protein phosphatase 1 (PP1)^25^. RIF1 deficiency, or its lack of interaction with PP1, cause both the loss of the temporal control of replication and the aberrant distribution of replication foci, inducing an accumulation of cells displaying early-like S-phase patterns^17,26,27^. Our trained model successfully recapitulates this observation in *Rif1*-deficient cell lines.

Given that a supervised approach not only requires pattern appearances to have been previously characterised, but also necessitates a sufficiently large annotated dataset for training, we next explore the results obtained via an unsupervised approach, which we applied to the same tasks. Using a popular self-supervised learning framework named Bootstrap Your Own Latent (BYOL)^28^, we successfully identify the abnormal replication profile of *Rif1*-deficient cell lines, while additionally reconstructing S-phase progression. We further validate our pipeline by employing the same unsupervised framework to analyse DNA replication in a well-characterised model of deregulated activation of replication origins, specifically in cells where cyclin E can be inducibly overexpressed^29^. The analysis of EdU and PCNA foci yields comparable results, thereby potentially paving the way for studying DNA replication dynamics in human tissue sections.

## Results

### A ResNet-50 convolutional neural network accurately assigns nuclei to pre-defined S-phase stages

Manual image analysis of the spatial distribution of EdU-labelled replication foci has been an invaluable tool for studying DNA replication dynamics, leading to the initial discovery and characterisation of the deregulation of replication timing induced by *Rif1* deletion in different cell types, including mouse embryonic fibroblasts (MEFs)^17^ and mESCs^27^. We have therefore chosen this well-characterised model to build and test our pipeline. Here, we pulsed wild-type mESCs for 30 minutes with EdU and cytospun them onto slides, thus obtaining a homogeneous monolayer of well-separated individual cells. To generate a sufficiently large dataset for machine learning, we acquired images at 40x magnification using the Olympus ScanR High Content Screening Microscope.

Following segmentation and filtering (See ‘Methods’), we manually categorised S-phase nuclei, initially assigning them to 3 classes (early, mid and late), with the subsequent addition of two intermediate classes (Fig. 1a). We introduced the latter two classes to account for commonly detected mixed patterns, corresponding to the early-mid and mid-late transitions. 858 S-phase nuclei were manually categorised, with the following distribution (according to early, early-mid, mid, mid-late and late stages): 384, 210, 110, 100 and 54 (Fig. 1b). This manually categorised dataset was split with stratified sampling into training, validation and test set according to a 60:20:20 ratio.

**Figure 1.**
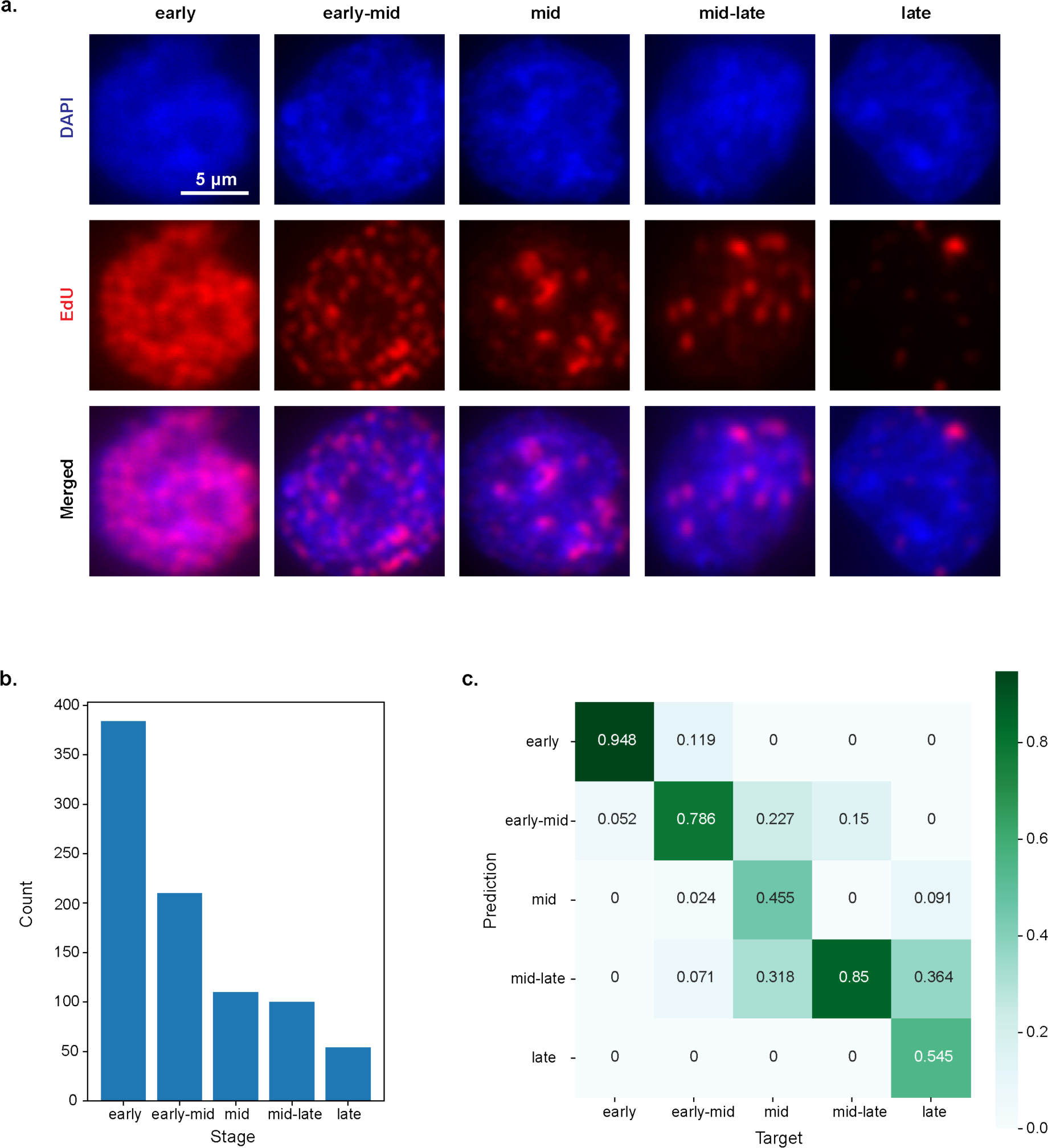
A ResNet-50 convolutional neural network accurately classifies nuclei into S-phase stages. **a.** Representative images of five S-phase categories, staged based on DAPI and EdU patterns. The criteria for manual categorisation are described in the ‘Methods’ section. **b.** Bar plot showing the distribution of 858 EdU-labelled nuclei. Nuclei count (from early to late) was as follows: 384, 210, 110, 100 and 54. **c.** Confusion matrix showing the performance of a pre-trained ResNet-50 convolutional neural network in classifying the different stages of S-phase. Classification performance was evaluated on the test set.

Using a ResNet-50 convolutional neural network^30^ pre-trained on ImageNet^31^, we achieved an overall classification accuracy of 80.8% ± 5.9% (95% CI) on the test set. Predictions were only considered correct when the exact target class was predicted. Altogether, we obtained reasonable prediction accuracies across all classes (Fig. 1c), considering that S-phase exists as a continuum of stages rather than a set of discrete states, with considerable overlap in observed patterns between adjacent classes. Henceforth, we refer to this trained model as ‘S-phase classifier’.

### A supervised learning model can reconstruct S-phase progression

During model training in an image classification setting, deeper layers in a neural network automatically learn a high-dimensional representation of each image, capturing high-level semantically useful features for distinguishing different target classes^32^. This representation, while potentially more informative than the class assigned to each image, cannot be directly visualised due to its high-dimensional nature. Here, we thus reduced the learned representations, or embeddings, encoded by the last hidden layer of the ‘S-phase classifier’ to two maximally informative dimensions using principal component analysis (PCA) to facilitate visualisation. This was performed for the manually categorised dataset comprising 858 wild-type S-phase nuclei.

PCA plots of the image embeddings revealed a clear gradient from early to late S-phase, showing that the ‘S-phase classifier’ had learned to organise images of S-phase nuclei in a sequence that recapitulates progression through S-phase (Fig. 2a). This is despite being provided no information regarding ordering amongst the classes. Although visualisation based on non-linear dimensionality reduction techniques such as t-SNE and UMAP has recently been challenged^33^, the corresponding analyses yielded results consistent with those based on PCA (Fig. 2b and 2c).

**Figure 2.**
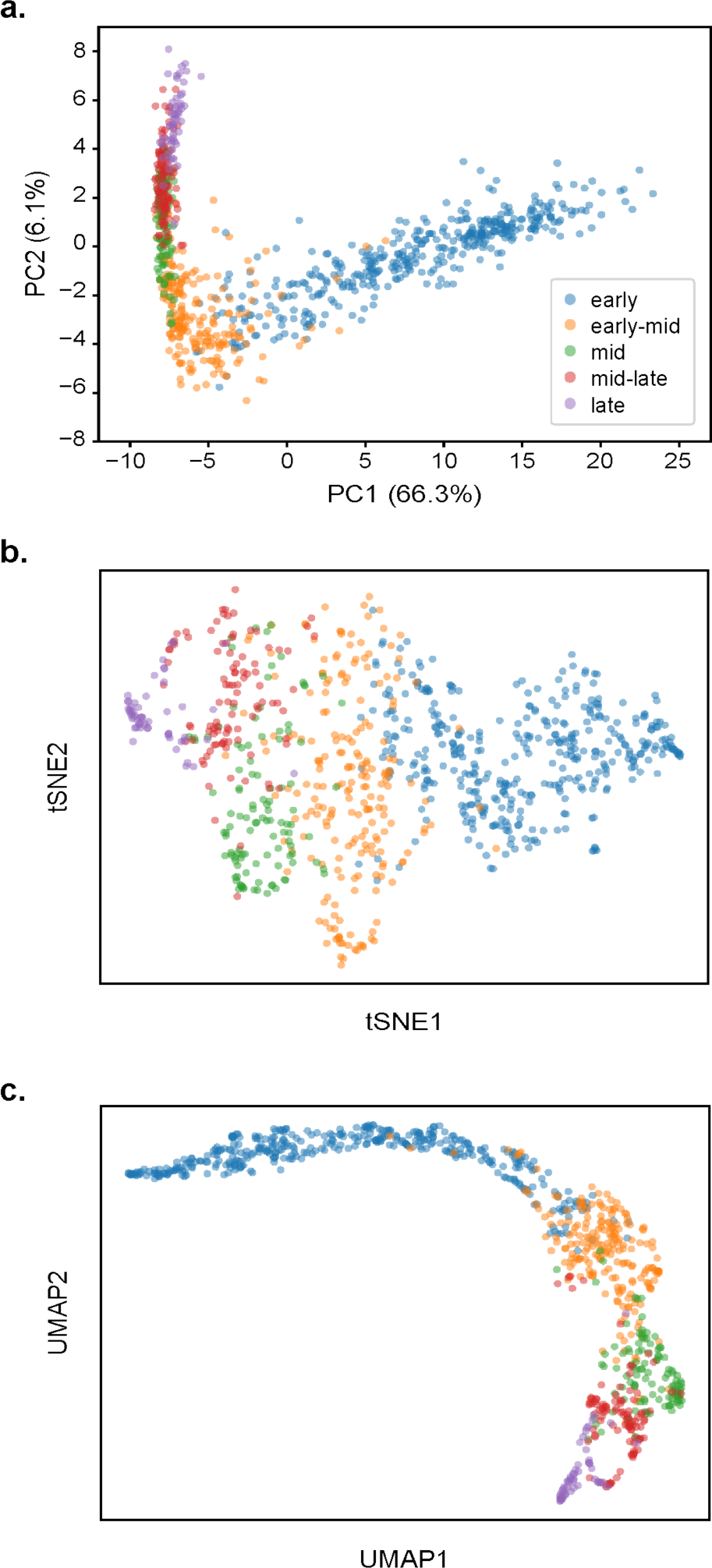
A supervised learning model reconstructs S-phase progression based on categorical labels. **a.** Scatter plot of the PCA-projected image embeddings encoded by the ‘S-phase classifier’ for the manually categorised dataset comprising 858 S-phase nuclei. A clear gradient from early S-phase (blue) to late S-phase (purple) can be observed despite no ordering information being provided to the model. Scatter plots based on tSNE and UMAP are, respectively, shown in **b.** and **c.** The plots are coloured according to the class predicted by the ‘S-phase classifier’.

### A supervised learning model can identify aberrant DNA replication dynamics

We have previously shown that expression of a *Rif1* allele carrying a point mutation that abolishes the interaction with PP1 (*Rif1^ΔPP1/-^)*, as well as *Rif1* deficiency (*Rif1^-/-^*), result in an increased proportion of early-like S-phase patterns, which parallels the deregulation of the order of firing of replication origins^26^. Although *Rif1* haploinsufficiency (*Rif1^TgWT/-^*) also affects the spatial distribution of EdU-labelled replication foci (albeit to a lesser extent), this is not accompanied by a change in replication timing^26^. Here, we have used *Rif1* wild-type and mutant mESCs to assess the performance of the ‘S-phase classifier’.

Following treatment with 4-hydroxytamoxifen (OHT)—which induces the conversion of the *Rif1*-flox allele (*Rif1^flox^*) into a *Rif1*-null allele (*Rif1^-^*)—a marked increase in the proportion of nuclei predicted as ‘early’ is observed for *Rif1^ΔPP1/-^* and *Rif1^-/-^* (Fig. 3a), successfully recapitulating previous results obtained by manual scoring^26^. Intriguingly, kernel density estimate (KDE) plots of the projected embeddings further revealed that early S-phase nuclei followed a more ‘compact’ distribution in *Rif1^ΔPP1/-^* and *Rif1^-/-^* (Fig. 3b; Supplementary Fig. 1a and 1b). Indeed, this was also observed for *Rif1^TgWT/-^*, despite no increase in nuclei being predicted as ‘early’ following OHT treatment. This suggests that the ‘S-phase classifier’, in addition to performing comparably to human operators when assigning S-phase stages, could also allow other aspects of aberrant DNA replication to be identified through visualisation of its encoded embeddings.

**Figure 3.**
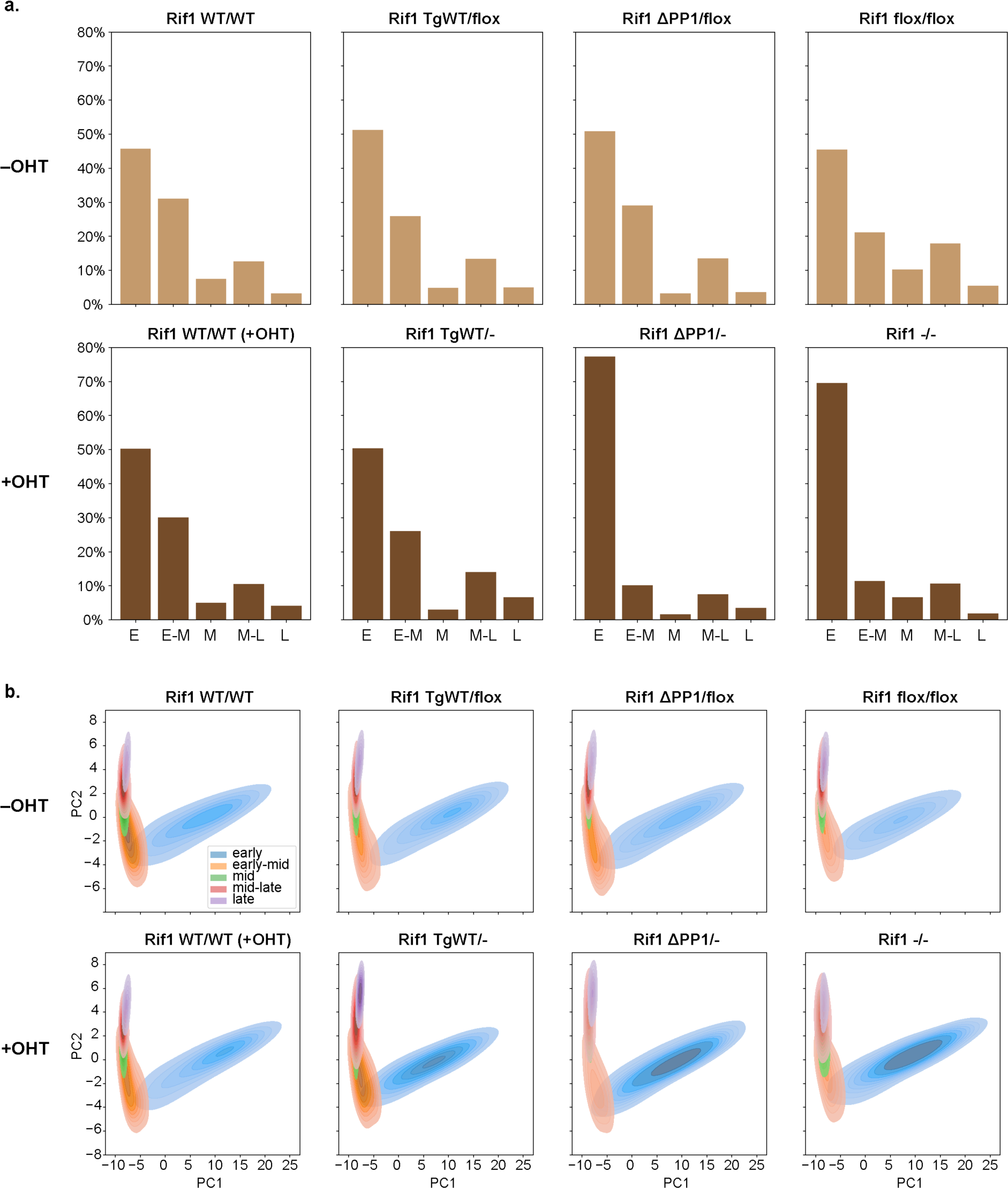
A supervised learning model identifies aberrant DNA replication dynamics upon Rif1 loss of function. **a.** Bar plots showing, per genotype, the proportion of nuclei assigned by the ‘S-phase classifier’ to each of the five categories: early, early-mid, mid, mid-late and late. The difference in distribution following addition of OHT was statistically interrogated based on Chi-squared tests, yielding the following results: Rif1*^WT/WT^* (+OHT): *P*= 1.4 x 10^-14^; Rif1*^TgWT/-^*: *P*= 9.6 x 10^-4^; Rif1*^ΔPP1/-^*: *P*= 9.7 x 10^-168^; Rif1*^-/-^*: *P*= 8.4 x 10^-90^. The top and bottom row: pre- and post-OHT treatment. **b.** KDE plots of the projected image embeddings encoded by the ‘S-phase classifier’ for each genotype, showing changes in distribution following OHT treatment. The plots are coloured according to the class predicted by the ‘S-phase classifier’.

### An unsupervised learning model can autonomously reconstruct S-phase progression and identify aberrant DNA replication dynamics

The embeddings learned in a supervised setting are determined by pre-defined categories. In this case, the different observed patterns in S-phase have been discriminated based on human assessment. Therefore, we sought to assess how the previous embeddings would compare to those learned using an unsupervised approach. Self-supervised learning is one increasingly popular such framework and has found considerable success in the field of computer vision^34–36^. Using automatically generated labels from unannotated data, approaches based on this framework aim to learn representations that allow generalisation to downstream tasks. Given that self-supervised image representation learning techniques could allow for complex patterns to be automatically extracted across a large dataset—while avoiding the need for manual annotation—they are an appealing prospect for discovery. Here, we employ a recent related framework, known as BYOL^28^.

After training the BYOL model on an uncategorised set of wild-type nuclei, we visualised the image embeddings for the full dataset comprising 45,128 S-phase nuclei, as performed previously for the manually categorised dataset. Even with no information regarding categorisation being provided to the model, the latter was still able to recapitulate progression through S-phase (Fig. 4a: right). This progression is more obvious when a densely clustered region is expanded, showing a gradient from early to late S-phase in the negative PC1 direction (Fig. 4a: left).

**Figure 4.**
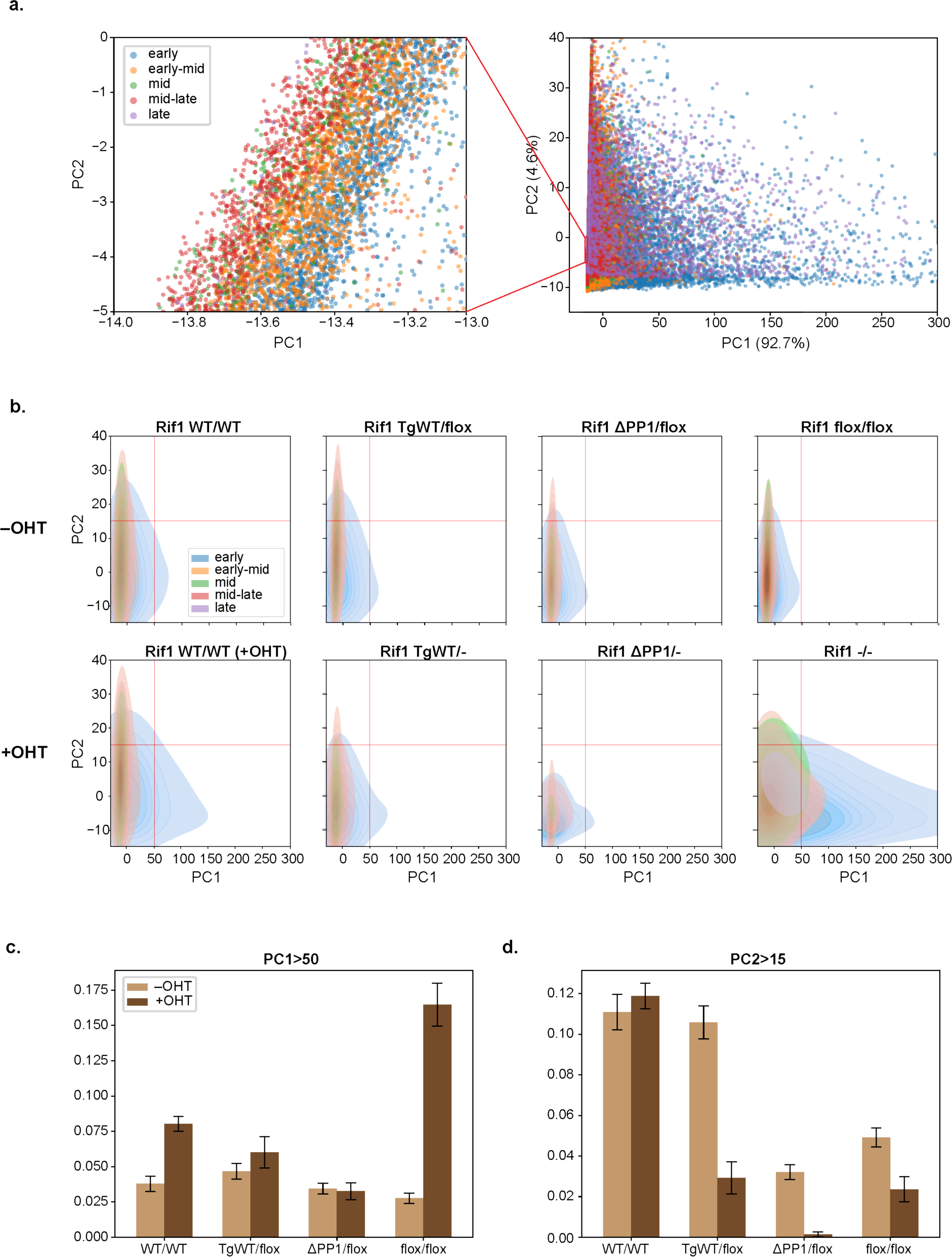
An unsupervised learning model reconstructs S-phase progression and identifies aberrant DNA replication dynamics upon *Rif1* loss of function. **a.** Scatter plot of the projected image embeddings encoded by the BYOL-trained model for the full dataset comprising 45,128 S-phase nuclei. A gradient from early S-phase (blue) to late S-phase (purple) can be observed. Axes were limited due to the presence of outliers, excluding 177 nuclei in total (145 nuclei with PC1 value greater than 300 and 32 nuclei with PC2 value greater than 40). **b.** KDE plots of the projected image embeddings encoded by the BYOL-trained model for each genotype, showing changes in distribution following OHT treatment. The plots in **a.** and **b.** are coloured according to the class predicted by the ‘S-phase classifier’. **c.** and **d.** Bar plots showing the proportion of nuclei, per genotype, with **c.** PC1 values greater than 50 and **d.** PC2 values greater than 15. The error bars represent 95% confidence intervals. The difference in proportion following addition of OHT was statistically interrogated based on two-proportion Z-tests, yielding the following results for **c**: Rif1*^WT/WT^*: *P*=8.2 x 10^-23^; Rif1*^TgWT/flox^*: *P*= 0.024; Rif1*^ΔPP1/flox^*: *P*= 0.59; Rif1*^flox/flox^*: *P*= 1.4 x 10^-134^ and for **d**: Rif1*^WT/WT^*: *P*= 0.15; Rif1*^TgWT/flox^*: *P*= 5.2 x 10^-23^; Rif1*^ΔPP1/flox^*: *P*= 3.0 x 10^-23^; Rif1*^flox/flox^*: *P*= 1.4 x 10^-7^.

KDE plots of the projected embeddings revealed marked changes in the distribution of *Rif1^ΔPP1/-^* and *Rif1^-/-^* (Fig. 4b; Supplementary Fig. 2a and 2b)—we have attempted to quantify the changes resulting from OHT treatment in Fig. 4c and 4d to aid interpretation, although we acknowledge that results based on separate analyses of PC1 and PC2 may not fully reflect the overall shift in distribution, which are otherwise only qualitatively assessed here. Interestingly, as opposed to the ‘S-phase classifier’, the BYOL-trained model detected distinct alterations in *Rif1^ΔPP1/-^* and *Rif1^-/-^*, with PC2 changes in both genotypes, but an increase in PC1 only in *Rif1^-/-^* cells. This parallels known biological differences: although both *Rif1* absence and loss of interaction with PP1 cause the loss of RT and an increase in early-like S-phase patterns, only *Rif1* absence is accompanied by an increase in the number of replication foci^26^. The BYOL-trained model additionally identified a change in PC2 in *Rif1^TgWT/-^* (Fig. 4b; Supplementary Fig. 2a and 2b), where only an increase in early-like S-phase patterns, without a change in the number of replication foci or in the RT program, has been documented^26^. It is therefore tempting to speculate that PC2 could reflect the alteration in the spatial distribution of replication foci, while PC1 could be related to the number of replication foci. Following OHT treatment, a shift in PC1 distribution is also observed in *Rif1^WT/WT^* (+OHT), though of a much smaller magnitude than in *Rif1^-/-^*. This suggests a possible effect on DNA replication due to the DNA damage response caused by induction of the Cre recombinase^37^. Overall, these findings support the idea that an unsupervised approach may be more sensitive than manual inspection at capturing various alterations in DNA replication.

### Unsupervised learning applied to the study of oncogene-induced deregulation of DNA replication dynamics

As a first step towards applying our approach to patient samples, we sought to extend our unsupervised framework to a different cellular system, closer to known pathological processes. For this purpose, we used a U2OS TetON CycE cell line^29^ (See ‘Methods’ for details). The overexpression of the oncogene cyclin E1 induces the precocious firing of an excessive number of origins, located within gene bodies, thus causing nucleotide depletion^38^ and replication fork collapse^20^. The rapid induction afforded by the TetOn system allowed us to monitor the early events that accompany aberrant origin induction. As expected, doxycycline treatment caused the rapid induction of cyclin E1 expression (Supplementary Fig. 3a) and the consequent increase in the proportion of cells entering S-phase (Supplementary Fig. 3b). By 96 hours, the proportion of cells in S-phase declines, the DNA damage checkpoint having been engaged^29^. This time, we visualised replication foci both by EdU labelling and by performing immunostaining for PCNA (Supplementary Fig. 4a). We analysed the images using an unsupervised approach analogous to the one described above for *Rif1* mutant mESCs.

After training the BYOL model on images of non-induced U2OS cells (‘0h’ timepoint), we proceeded to visualise the image embeddings for the entire dataset, consisting of 26,137 S-phase nuclei across the five timepoints (Supplementary Fig. 4b). Two separate models were trained: the first on images consisting of the DAPI and EdU channels (DAPI/EdU) and the second based on the DAPI and PCNA channels (DAPI/PCNA). In both cases, KDE plots showed a gradual change in the overall distribution from the ‘0h’ to the ‘48h’ timepoint, with a shift towards increasing PC1 and PC2 values for the DAPI/EdU embeddings (Fig. 5a; Supplementary Fig. 5a), and respective negative and positive shifts in the distribution of PC1 and PC2 for the DAPI/PCNA embeddings (Fig. 5b; Supplementary Fig. 5b). From the ‘48h’ to the ‘96h’ timepoint, a shift in the opposite direction is then observed for both embeddings. We assess these changes for the DAPI/EdU embeddings in Fig. 5c, and for the DAPI/PCNA embeddings, in Fig. 5d and 5e. Altogether, while our approach again does not allow us to separately explain the changes in PC1 and PC2, we find that the overall patterns in the learned embeddings reflect known biology. These results demonstrate that high-throughput image acquisition of PCNA-immunostained human cells can be successfully combined with unsupervised deep learning to automatically detect DNA replication aberration that may otherwise be challenging to reproduce by manual scoring.

**Figure 5.**
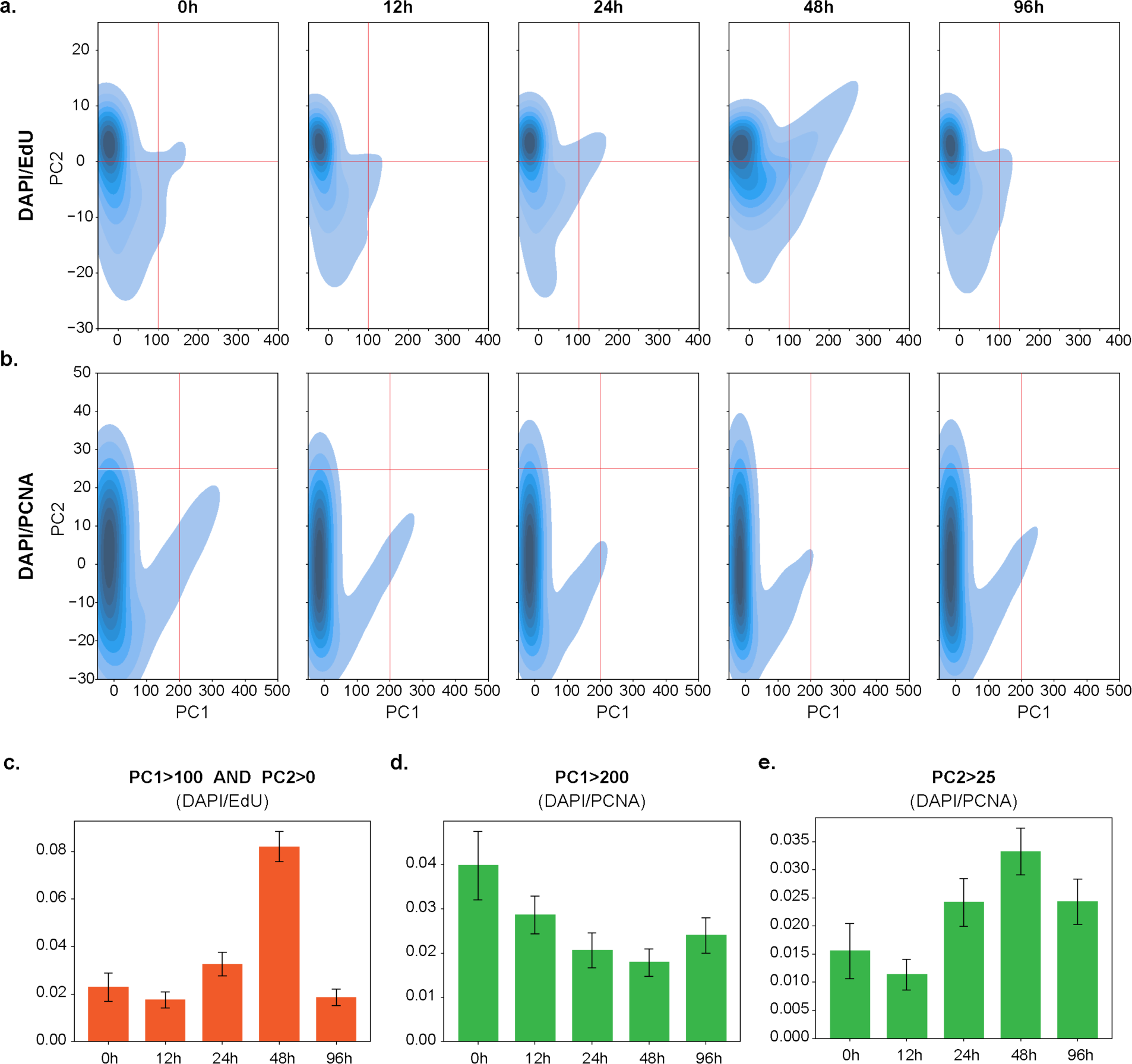
EdU- or PCNA-based analyses both identify the progressive deregulation of origin firing upon cyclin E1 induction. KDE plots of the projected image embeddings encoded by the **a.** DAPI/EdU **b.** DAPI/PCNA model. A progressive change in distribution is seen for both embeddings from the ‘0h’ to the ‘48h’ timepoint, with a shift in the opposite direction being observed from the ‘48h’ to the ‘96h’ timepoint. For the DAPI/EdU embeddings, PC1 and PC2 explained 98.2% and 1.2% of the variance, respectively. For the DAPI/PCNA embeddings, PC1 and PC2 explained 93.7% and 5.4% of the variance, respectively. **c.** Bar plots showing the proportion of nuclei, per timepoint, with both PC1 values greater than 100 and PC2 values greater than 0 (based on the DAPI/EdU embeddings). P-values for differences in proportion following addition of doxycycline are as follows: 12h: *P*= 0.11; 24h: *P*= 0.020; 48h: *P*= 6.2 x 10^-24^; 96h: *P*= 0.21. **d.** and **e.** Bar plots showing the proportion of nuclei, per timepoint, with **d.** PC1 values greater than 200 and **e.** PC2 values greater than 25 (based on the DAPI/PCNA embeddings). P-values for differences in proportion following addition of doxycycline are as follows for **d**: 12h: *P*= 8.5 x 10^-3^; 24h: *P*= 1.7 x 10^-6^; 48h: *P*= 7.7 x 10^-10^; 96h: *P*= 9.8 x 10^-5^ and for **e**: 12h: *P*= 0.12; 24h: *P*= 0.015; 48h: *P*= 6.5 x 10^-6^ ; 96h: *P*= 0.013. Differences in proportion between the ‘0h’ timepoint and other timepoints were assessed using two-proportion Z-tests. The error bars in **c.**, **d.** and **e.** represent 95% confidence intervals.

## Discussion

With deep learning-based approaches becoming increasingly popular for image analysis in the biological sciences^39^, we take advantage here of well-characterised biological systems to demonstrate how both supervised and unsupervised learning techniques can be applied to the study of DNA replication. Over the past two decades, genomic techniques have significantly advanced our understanding of DNA replication regulation. However, these approaches are costly and require specialised expertise for both sample preparation and analysis. Moreover, genomic methods require large numbers of cells and often rely on labelling replicating DNA with thymidine analogues, making them unsuitable for studying patient samples. Currently, analysing replication foci represents the most straightforward and cost-effective approach to study DNA replication, providing simultaneous information on the temporal and spatial control of replication. However, manual categorisation of images requires training, time and is susceptible to biases. Additionally, manual inspection alone may fail to extract complex patterns contained in a dataset, which could limit discovery potential compared to machine learning-based approaches.

Here, we first show that a convolutional neural network can classify with reasonable accuracies the different S-phase stages in fluorescence microscopy images of DAPI-stained and EdU-pulsed mESCs. By leveraging the trained model to undertake large-scale analysis, we successfully identify the altered DNA replication dynamics associated with *Rif1*-deficient mESCs, revealing a skewed distribution across the various S-phase stages, consistent with our previous findings. Nonetheless, although employing such a model trained through supervised learning enables rapid and consistent categorisation of images, supervised techniques still require access to a sufficiently large and representative manually annotated dataset for model training. More generally, supervised learning techniques rely on the appearance of cell type-specific patterns having been previously characterised, and are therefore not easily applied when image annotation is challenging or impractical, or has simply not been attempted.

Against this background, unsupervised approaches are particularly relevant, necessitating no label input and therefore easily generalising to such scenarios. Moreover, the embeddings encoded by an unsupervised learning model reflect inherent structure in the data, rather than that imposed through human assessment. Incorporating such techniques as part of high-throughput experiments is thus a potentially fruitful avenue. Importantly, approaches similar to ours—which may be considered as a form of ‘image-based profiling’^40,41^—could also be applied to other settings where distinct patterns, not necessarily based on EdU labelling, reflect different underlying cellular states. More widespread adoption will require the generation of large annotated datasets and associated benchmarks^42^ to act as a test bed for evaluating the robustness and generalisation potential of different deep learning techniques, of which we have selected only one specific example here. Alongside this, continued advances in deep learning that aid model interpretability will be crucial.

Finally, we have demonstrated that PCNA and EdU provide equivalent information, thus potentially allowing for high-throughput imaging analysis of DNA replication to be extended to human sample studies in the future. Indeed, most of our knowledge pertaining to the involvement of aberrant DNA replication in pathogenesis comes from developmental diseases^18^. Very little is known, except for cancer^38,43,44^, regarding whether and how somatic mutations in any of the DNA replication-controlling genes contribute to disease. Yet, the known association between inherited mutations and, for example, immunological dysfunctions suggests that investigating the role of deregulated DNA replication in sporadic pathologies could be an important pursuit, which will be greatly facilitated by approaches, such as ours, based on high-throughput imaging.

## Methods

### Cell lines

The mESC cell lines used in this dataset comprised four genotypes, with two clones per genotype included as biological replicates. The four genotypes are as follows: *Rif1^WT/WT^*, *Rif1^TgWT/flox^*, *Rif1^ΔPP1/flox^* and *Rif1^flox/flox^*, all in combination with *Rosa26^Cre-ERT/+^*. For *Rif1^flox/flox^*, see ref.^27,37^; for a detailed description of *Rif1^TgWT/flox^* and *Rif1^ΔPP1/flox^*, see ref.^26^. Briefly, in *Rif1^TgWT/flox^* and *Rif1^ΔPP1/flox^* mESCs, the *Rif1* conditional allele (*Rif1^flox^*) carries loxP sites flanking exons 5 and 7; the second *Rif1* allele contains a knock-in of either a *Rif1* wild-type mini-gene (*TgWT*) or a mini-gene encoding *Rif1^ΔPP1^*, where PP1 interaction sites have been mutated. Addition of OHT induces Cre-loxP recombination, which abolishes RIF1 expression from *Rif1^flox^* (converting it to *Rif1^-^*), leaving either *Rif1^ΔPP1^* or *Rif1^TgWT^* as the only source of RIF1 (hemizygosis). In the absence of OHT, all cell lines are phenotypically wild-type. For information regarding derivation and culture and experimental conditions, see ref.^26^.

The U2OS TetON CycE cell line was kindly gifted by the Difley Lab^29^. U2OS cells were cultured at 37°C in 5% CO_2_ using DMEM (Gibco, 41965-062) supplemented with 10% heat-inactivated Tet-free fetal bovine serum and 1% penicillin/streptomycin (Gibco 15070063). Overexpression of cyclin E1 was induced using 1 μg/ml doxycycline (Sigma Aldrich, Cat# D9891).

### Immunofluorescence and Western blot

mESCs were pulsed with 10 μM EdU for 30 minutes. Harvested cells were resuspended in 1% FBS/PBS. 60,000 cells per sample were cytospun to poly-lysine-coated slides by centrifugation at 800 rpm, followed by fixation with 4% PFA for 10 minutes at room temperature. After PBS washes, samples were permeabilised with 0.5% NP40/PBS for 10 minutes and washed with PBS. Samples were then incubated with blocking buffer (1% BSA in TBST) for 30 minutes at room temperature. Click-iT reaction was performed according to the manufacturer’s instruction (Invitrogen, C10424), using Alexa Fluor 647 (Invitrogen, A100277). Samples were then stained with DAPI at 2.5 μg/ml in PBS for 30 minutes. After washes, in-house mounting medium (0.1% w/v, p-phenylenediamine; 90% v/v, glycerol in PBS) was used to mount coverslips. Images were taken with an Olympus ScanR High Content Screening Microscope, using a 40x objective. Of note, although adopting the cytospin preparation method greatly benefitted our approach—ensuring adequate separation between cells and thus greatly facilitating the task of segmentation—its high-speed nature resulted in a subset of nuclei with abnormal morphology.

U2OS cells, in contrast to mESCs, grow as a monolayer of well-separated individual cells and were therefore directly grown on gelatinised coverslips. U2OS cells were pulsed with 10μM EdU for 30 minutes. After PBS washes, cells were pre-extracted with Triton buffer (0.5% Triton X-100; 20 mM Hepes-KOH (pH 7.9); 50 mM NaCl; 3 mM MgCl2; 300 mM Sucrose) for 5 minutes at 4°C, before fixation with ice-cold methanol for 7 minutes at 4°C. After PBS washes, samples were permeabilised with Triton buffer for 10 minutes at room temperature and washed with PBS. Samples were then incubated with PBG blocking buffer (0.2% w/v, cold water fish gelatin-Sigma G-7765; 0.5% w/v, BSA-Sigma A-2153 in PBS) for 30 minutes at room temperature. The primary antibody against PCNA (Abcam, ab29 [PC10]) was diluted 1:1000 in blocking solution, and then applied to samples for a 2-hour incubation at room temperature. Following washes with blocking buffer, samples were subsequently incubated with both the secondary antibody (Invitrogen, diluted 1:800) and DAPI (2.5 μg/ml) for 45 minutes in dark conditions. The immunofluorescence signals were fixed with 4% PFA for 10 minutes. Click-iT reaction was performed as described above, after which the coverslips were mounted with Vectashield (Vector Laboratories). SDS-PAGE and western blot analysis were performed as per ref.^27^. The following antibodies were used for western blotting: anti-HA (Covance monoclonal HA.11 clone 16B12 #MMS-101R) 1:1000; and anti-lamin B1 (Abcam #ab16048) 1:2000.

### Segmentation and filtering

Following image acquisition, we employed a simple segmentation approach based on DAPI signal^45^, thresholding using Otsu’s method^46^, before extracting individual nuclei using the watershed algorithm^47^. Nuclei were filtered according to morphology, with additional thresholds based on image sharpness and minimum EdU intensity subsequently being applied (See below for inclusion criteria). 1,202 nuclei were retained after filtering, and upon manual inspection were found to include 143 G1/G2 nuclei, 140 nuclei with abnormal morphology, 49 blurry images and 12 inaccurately segmented images (Supplementary Fig. 6a). The remaining 858 S-phase nuclei were deemed suitable for manual categorisation. Given that manual inspection to identify S-phase nuclei was impractical for the full dataset, we decided to train a neural network on the manually categorised dataset comprising 1,202 nuclei. This ‘overall classifier’ distinguished between S-phase and G1/G2-phase with high accuracy (Supplementary Fig. 6b). The number of S-phase nuclei per genotype identified by the ‘overall classifier’ is given in Supplementary Fig. 6c.

For U2OS cells, segmentation and filtering steps were analogously performed, with morphology-related thresholds being adapted to U2OS cells. Thresholds for identifying S-phase nuclei were this time based on both the EdU and PCNA channels to allow comparison between the two markers (See below for inclusion criteria). Importantly, we opted here to apply more stringent filtering criteria to refine the dataset quality, instead of employing an ‘overall classifier’, which we acknowledge would inevitably limit the practical application of our pipeline. 26,137 S-phase nuclei were retained across the five timepoints (Supplementary Fig. 4b).

### Inclusion criteria

For mESCs, inclusion criteria with respect to morphology were as follows: minimum area of 2,000, maximum area of 10,000, minimum minor axis length of 40, maximum major axis length of 120, and maximum eccentricity of 0.7. Measurements provided here are in terms of pixels. For reference, each pixel corresponds to 0.1625 micron. A minimum mean EdU intensity of 1,000 was applied to limit the number of G1/G2 nuclei, but not excluding S-phase nuclei with low EdU intensity. To filter out images with high EdU intensity arising from neighbouring signal, we also set a minimum mean EdU intensity of 1000 for a 10×10 pixel square centred on the image centroid. After trialling various measures of sharpness, we used the blur metric implemented in ‘measure.blur_effect’ from the scikit-image package, using default parameters^48^. Images were filtered out if the estimated blur values on the DAPI channel or EdU channel were more than 0.75 or 0.7, respectively. All thresholds were decided after plotting the distribution histograms and manually inspecting nuclei corresponding to the distribution tails.

For U2OS cells, inclusion criteria with respect to morphology were as follows: minimum area of 2,000, maximum area of 25,000, minimum minor axis length of 40, maximum major axis length of 250, and maximum eccentricity of 0.8. A minimum mean EdU intensity of 2,000 and a minimum mean PCNA intensity of 500 were applied, with respective minimum mean EdU and PCNA intensities of 2000 and 500 similarly being set for a 10×10 pixel square centred on the image centroid. Images were filtered out if the estimated blur values on the DAPI channel, EdU channel or PCNA channel were more than 0.75, 0.7 or 0.7, respectively. We computed the mean Euclidean norm of the gradient of the image as an additional measure of sharpness, and excluded those images with sharpness values less than 100 and 50 on the EdU and PCNA channels, respectively. Finally, we filtered out images with high background EdU signal using the ‘exposure.is_low_contrast’ function from the scikit-image package, using default parameters.

### Manual categorisation

858 wild-type S-phase nuclei (*Rif1^TgWT/flox^*) were manually categorised. Nuclei with diffuse EdU signal were categorised as ‘early’. Nuclei with EdU-positive foci coinciding with chromocenters and those demonstrating peripheral EdU signal were categorised as ‘mid’, while those for which EdU-positive foci did not coincide with chromocenters were categorised as ‘late’. When the EdU signal involved features associated with both early and mid S-phase, or with both mid and late S-phase, nuclei were categorised as ‘early-mid’ and ‘mid-late’, respectively. In cases where no visible chromocenters were present in the DAPI channel and the EdU signal was foci-like in appearance, nuclei were categorised as ‘mid-late’. Manual categorisation for training the ‘overall classifier’ was performed according to the target classes (Supplementary Fig. 6a). Nuclei with poor morphology refer to those that are either damaged, distorted, broken or dead, acknowledging overlap between possible classes. Given the overlap between biological and machine learning terminology, we have reserved use of the term ‘labelled’ to refer to those nuclei that are EdU-positive, instead using the term ‘categorised’ when referring to nuclei that have been manually assigned to different classes.

### Supervised learning

A stratified split into training, validation and test set according to a 60:20:20 ratio was performed. Hyperparameters were fine-tuned on the validation set, with training and validation sets being subsequently combined. Results are reported on the test set. Datasets were normalised based on the training set per channel mean and standard deviation. Data augmentation was performed to artificially increase dataset size. It was limited here to random horizontal and vertical flips given that other augmentation techniques such as cropping or colour jitter may not preserve S-phase category, which is highly dependent on channel-specific information. Given the class imbalance in our dataset, which is skewed towards earlier S-phase stages, images were resampled from the training set at each epoch with sampling weights inversely proportional to the frequency of the class they belong to, i.e. images from more frequently occurring classes are sampled less often.

The ‘S-phase classifier’ and ‘overall classifier’ were both based on a ResNet-50 model architecture—a powerful 50-layer convolutional neural network with residual connections across different layers that has been successfully used in various computer vision tasks^30^. Although we are aware of more recent deep network architectures such as vision transformers^49^, we chose ResNet-50 due to its excellent performance/computation trade-off. The last fully-connected layer of the neural network was modified to have an output size corresponding to the number of target classes, and the model trained using a cross-entropy loss. Model training was performed using a batch size of 32 on a NVIDIA A100 GPU, with optimisation using stochastic gradient descent with a momentum of 0.9. A cosine annealing schedule was used with warm restarts every 20 epochs^50^. Initial and minimum learning rates were set to 1e-2 and 0, respectively. The model was trained until convergence using early stopping, with the best weights (based on validation accuracy) being returned upon training completion.

### Self-supervised learning

We used the BYOL implementation provided at the following GitHub repository: https://github.com/lucidrains/byol-pytorch (version 0.6.0). Parameters were set as follows: image_size = 224, hidden_layer = ‘avgpool’, projection_size = 256, projection_hidden_size = 4096, augment_fn = augment_fn, augment_fn2 = augment_fn2, moving_average_decay = 0.99, use_momentum = True. The model was trained using the Adam optimiser with a learning rate of 3e-4, and β1 and β2 values of 0.9 and 0.999, respectively. The first augmentation was set to random horizontal flips, while the second augmentation was set to random vertical flips, for reasons similar to those already described above. The model was trained from scratch instead of using ImageNet pre-trained weights, given the possibility here to train on a relatively large uncategorised dataset. The mESC training set for self-supervised learning included both clones of *Rif1^WT/WT^*, both clones of *Rif1^flox/flox^,* and the *Rif1^TgWT/flox^* clone not used for supervised learning, comprising 16,197 S-phase nuclei in total. The trained model was then used to generate embeddings for the full dataset consisting of 45,128 S-phase nuclei. The U2OS training set comprised the 2,443 S-phase nuclei corresponding to the ‘0h’ timepoint. Similarly, the trained model was then used to generate embeddings for the entire dataset consisting of 26,137 S-phase nuclei.

### Visualisation & Statistics

Following data centering, dimensionality reduction on the embeddings was performed using the implementation of PCA provided by the scikit-learn package. Image embeddings for individual genotypes were projected onto a common PC space (defined by the first two PCs derived from the full dataset) to facilitate comparison. KDE plots were generated via the ‘seaborn.kdeplot’ function, using default parameters except for the following: bw_adjust = 2 and thresh = 0.1. Chi-squared tests and two-proportion Z-tests were performed as implemented by the ‘scipy.stats’ and ‘statsmodels.stats.proportion’ modules, respectively, with uncorrected p-values being provided. 95% confidence intervals, which were based on a normal approximation, are given by:

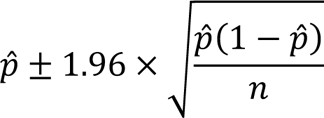

where 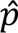 and *n* represent the sample proportion and sample size, respectively.

### Software

Scripts were written using Python version 3.7.12, with neural networks developed using PyTorch version 1.13.1 (CUDA version 11.7). Scientific analysis was carried out using NumPy version 1.21.6, pandas version 1.3.5, SciPy version 1.7.3 and scikit-learn version 1.0.2. Plots were made using Matplotlib 3.5.3 and Seaborn 0.12.1. Image analysis was performed using scikit-image version 0.19.3.

## Data availability

The image datasets associated with this paper will be made available upon publication.

## Code availability

The code for this project is available at the following GitLab repository: https://git.ecdf.ed.ac.uk/buonomo/deeplearning_dna_replication_2024

## Acknowledgements

We are grateful to the Diffley Lab including Jingkun Zeng and Lucy Drury for gifting the U2OS TetON CycE cell line to us. We also thank Matt Pearson and Ann Wheeler from the IGC Advanced Imaging Resource for their expertise & assistance regarding image acquisition. We would further like to acknowledge early work performed by Stefano Gnan and Eleonora Castelli that led to this project. This work made use of the resources provided by the Edinburgh Compute and Data Facility (ECDF) (http://www.ecdf.ed.ac.uk/).

## Author contributions

J.N.K.K. conducted the majority of the analyses and performed the experiments involving the U2OS TetON CycE cell line. B.P., F.N.C.S. and A.S. built the initial classification pipeline and provided feedback on the analyses. N.C. performed the experiments involving the *Rif1* mutant mESCs, with assistance on microscopy from C.A. S.C.B.B. conceived the project and jointly supervised the work with H.B. J.N.K.K. and S.C.B.B. wrote the manuscript with input from all authors.

## Supplementary

**Supplementary Figure 1.**
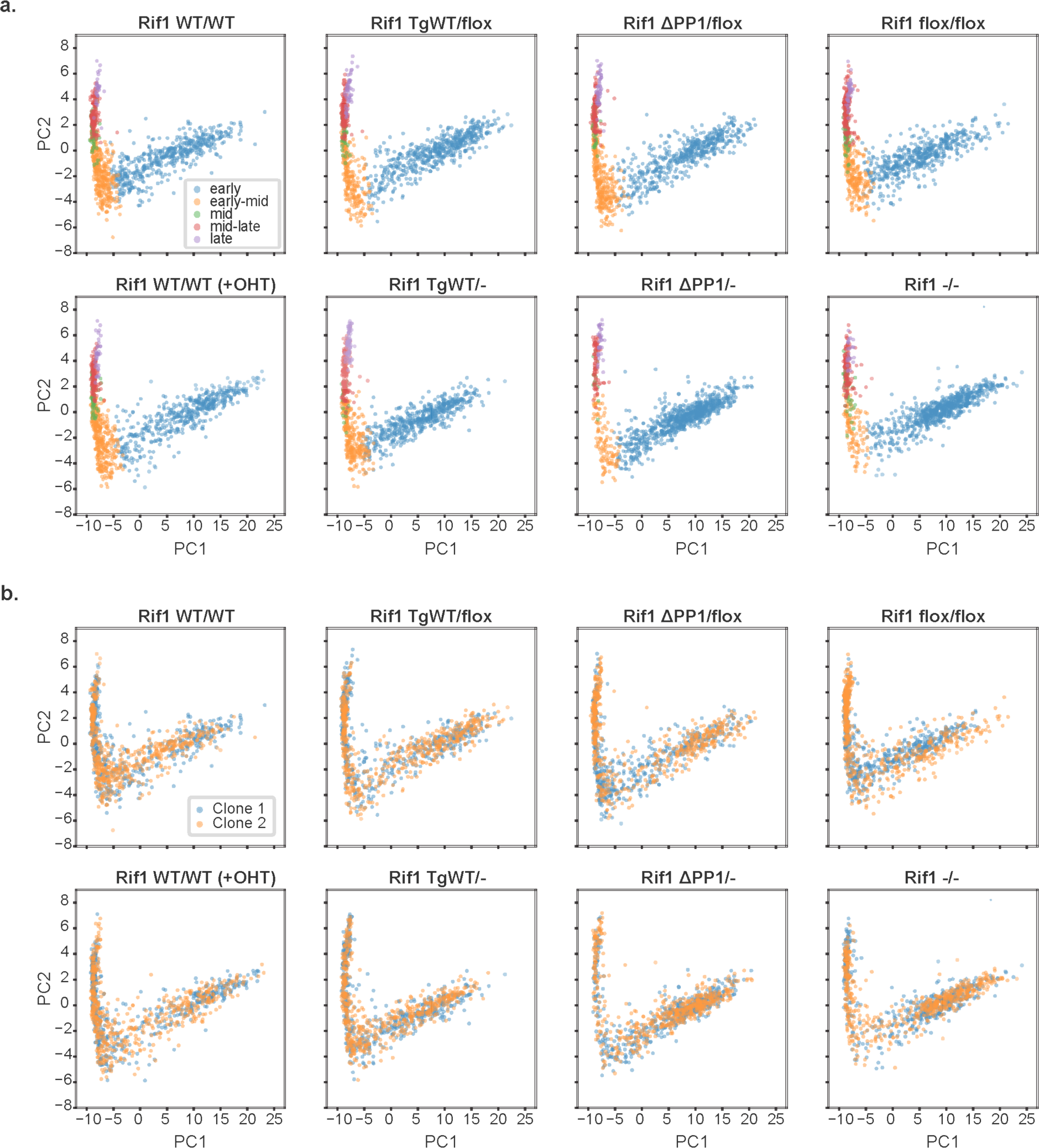
Aberrant DNA replication dynamics in *Rif1* mutant mESCs identified using a supervised approach. Scatter plots of the projected image embeddings, corresponding to the KDE plots in Fig. 3b. An equal number of nuclei per genotype (N = 1,000; 500 for each clone) was randomly sampled to facilitate comparison. The plots are coloured according to **a.** the class predicted by the ‘S-phase classifier’ and **b.** whether they belong to the first or second clone.

**Supplementary Figure 2.**
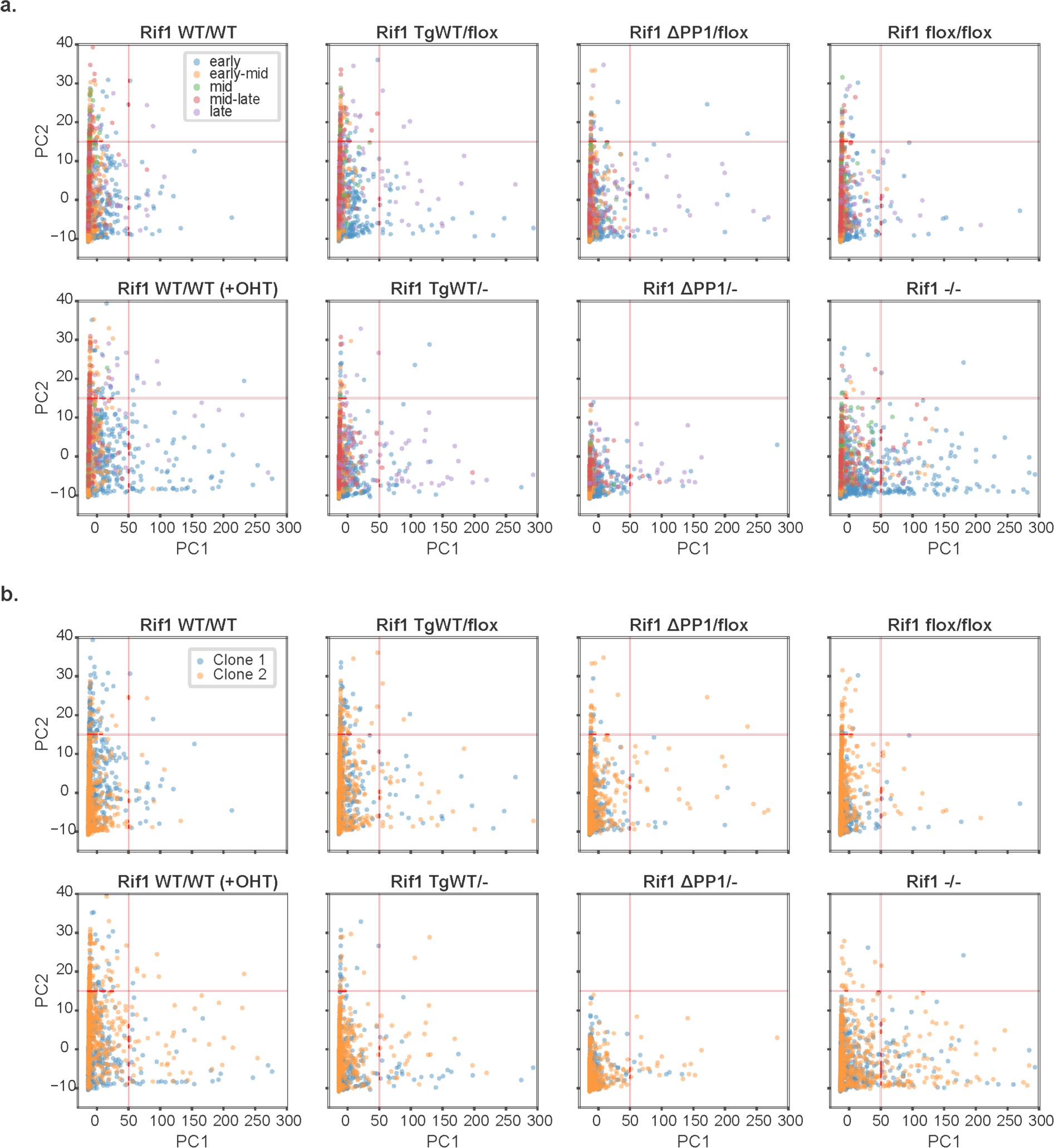
Aberrant DNA replication dynamics in *Rif1* mutant mESCs identified using an unsupervised approach. Scatter plots of the projected image embeddings, corresponding to the KDE plots in Fig. 4b. An equal number of nuclei per genotype (N = 1,000; 500 for each clone) was randomly sampled to facilitate comparison. The plots are coloured according to **a.** the class predicted by the ‘S-phase classifier’ and **b.** whether they belong to the first or second clone.

**Supplementary Figure 3.**
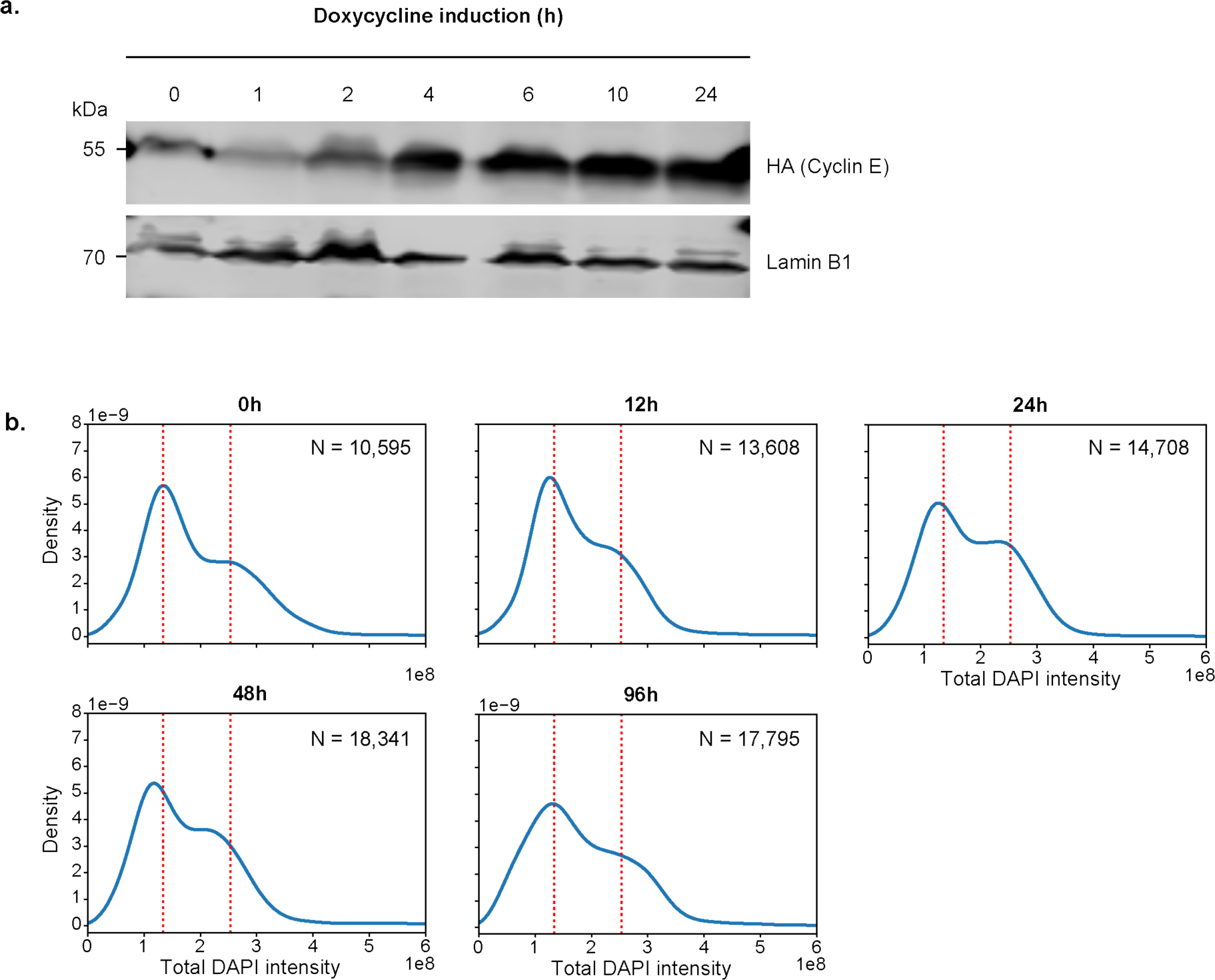
Doxycycline-induced expression of cyclin E1. **a.** Expression of cyclin E1-HA in U2OS TetON CycE cells was detected by immunoblots with anti HA-antibody. Lamin B1 was used as a loading control. **b.** KDE plots of the total DAPI intensity at each timepoint. The number of nuclei corresponding to each timepoint is shown. Vertical dotted lines are drawn in red at the approximate G1 and G2 positions (based on the ‘0h’ timepoint) to facilitate comparison across timepoints.

**Supplementary Figure 4.**
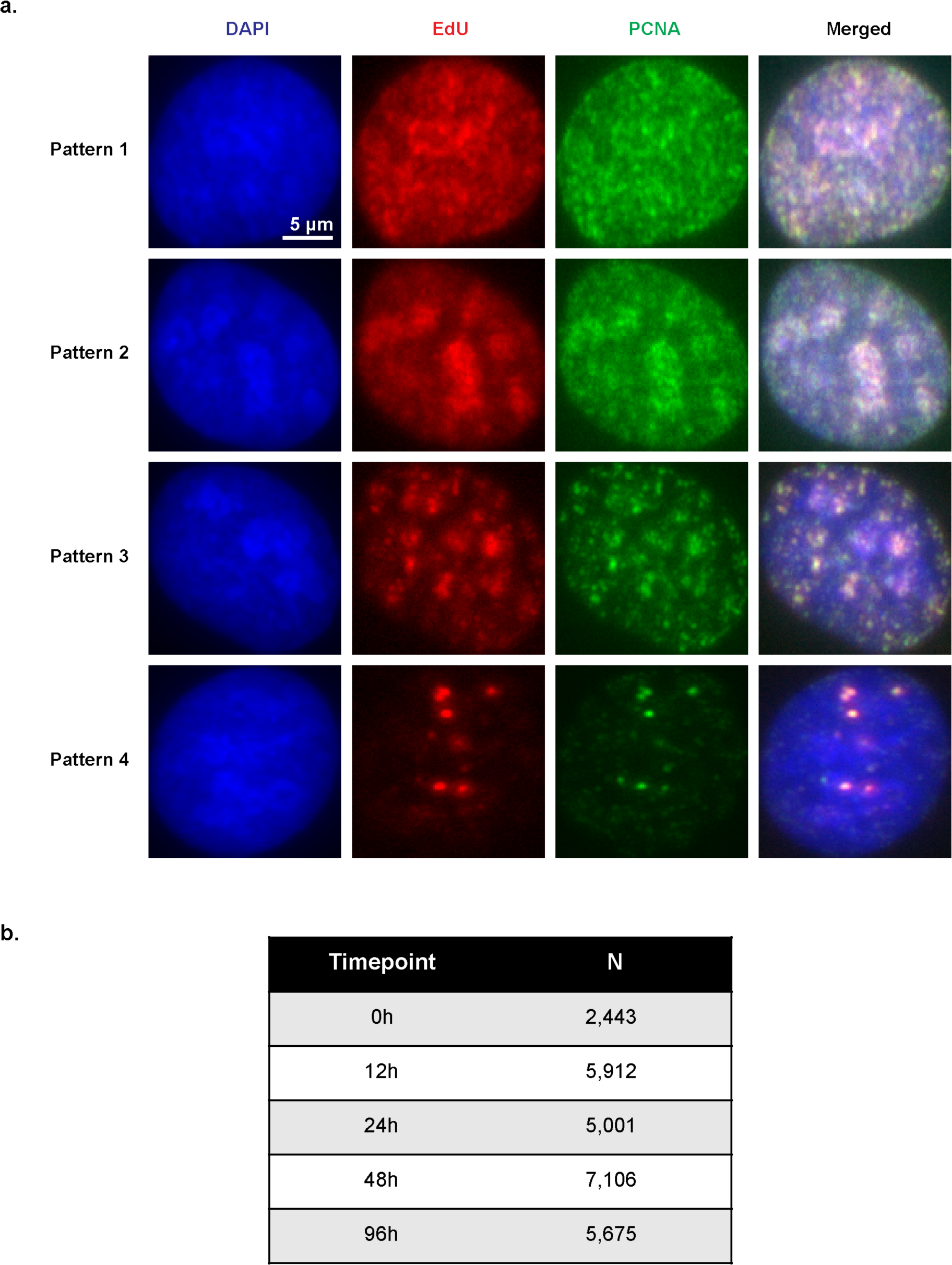
U2OS S-phase nuclei. **a.** Representative images of S-phase nuclei corresponding to four different observed patterns in U2OS cells (not treated with doxycycline) are shown. Images were taken with an Olympus ScanR High Content Screening Microscope, using a 40x objective. **b.** Table summarising the number of S-phase nuclei analysed per time point. In total, we identified 26,137 S-phase nuclei following segmentation and filtering steps.

**Supplementary Figure 5.**
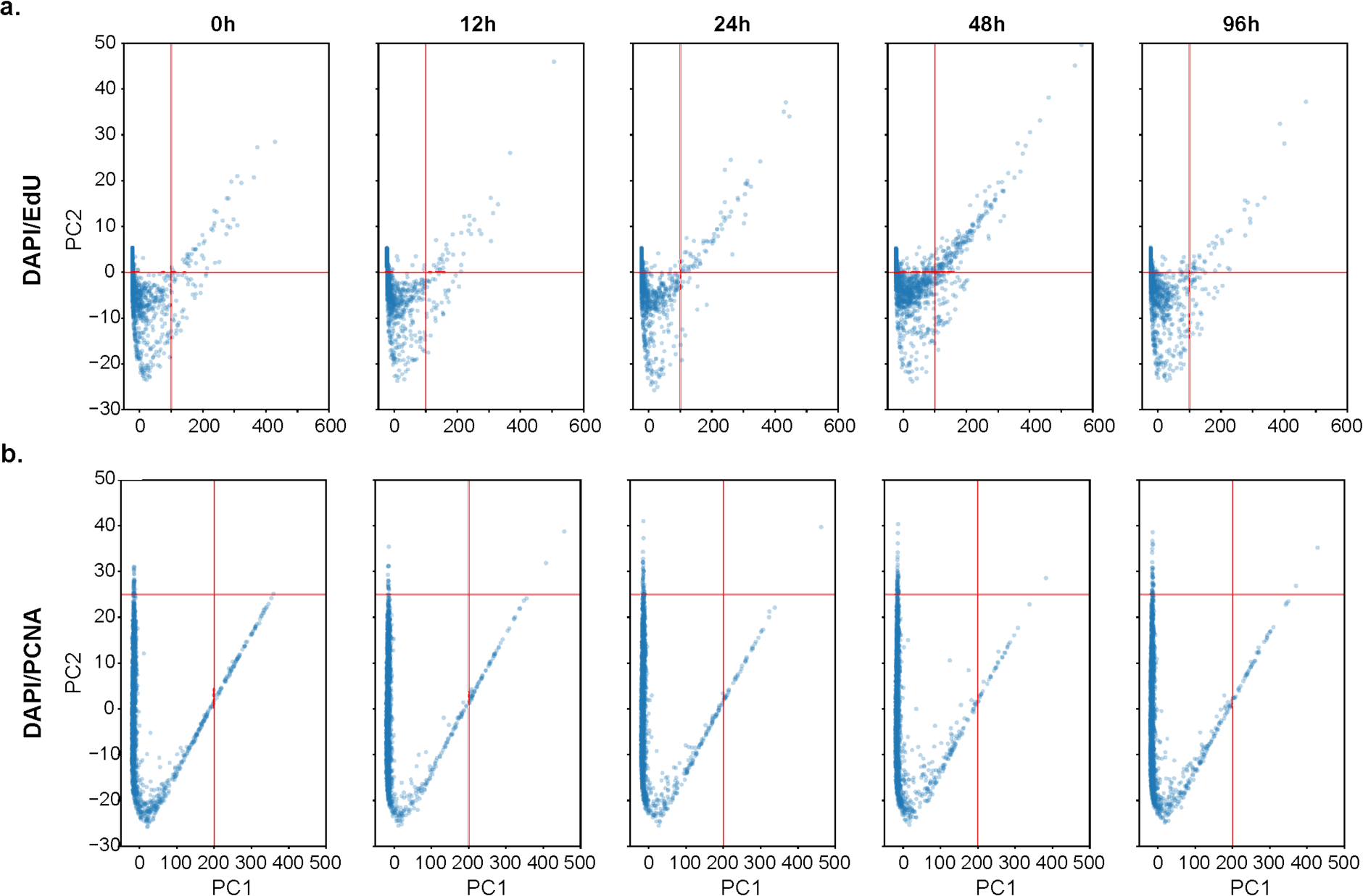
Deregulated origin firing in U2OS TetON CycE cells identified based on both EdU and PCNA. Scatter plots of the projected image embeddings, corresponding to the KDE plots in **a.** Fig. 5a. and **b.** Fig. 5b. An equal number of nuclei per timepoint (N = 2,000) was randomly sampled to facilitate comparison.

**Supplementary Figure 6.**
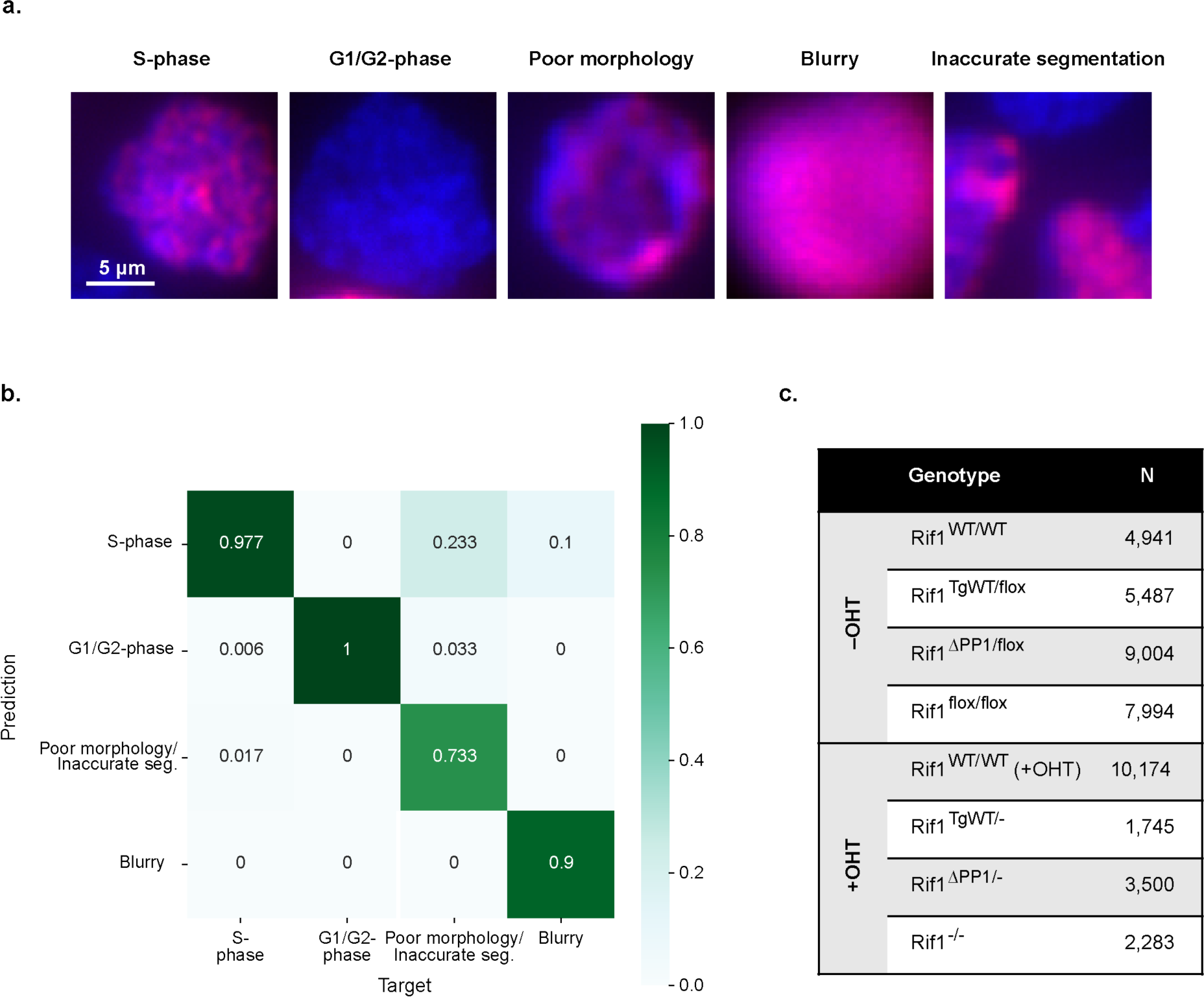
Use of a convolutional neural network (‘overall classifier’) to identify S-phase nuclei across the whole dataset. **a.** Representative images corresponding each of the five possible categories encountered during manual inspection of the 1,202 images retained following image segmentation and filtering. These 1,202 manually categorised images were used for training and testing a convolutional neural network (‘overall classifier’), whose aim was to identify S-phase nuclei across the whole dataset comprising both wild-type and mutant cell lines. The blue and red channels correspond to DAPI and EdU, respectively. **b.** Confusion matrix showing the performance of a pre-trained ResNet-50 convolutional neural network in identifying S-phase nuclei. Images of nuclei demonstrating poor morphology and those corresponding to inaccurate segmentation were considered as a single class given the small number of images present in the latter category. **c.** Table showing the number of S-phase nuclei, per genotype, present in our dataset. In total, we identified 45,128 S-phase nuclei across the full dataset following segmentation and filtering steps.

